## Supplementary Figures for "High-throughput interrogation of programmed ribosomal frameshifting in human cells"

This PDF file includes:

Supplementary Figures S1 to S12

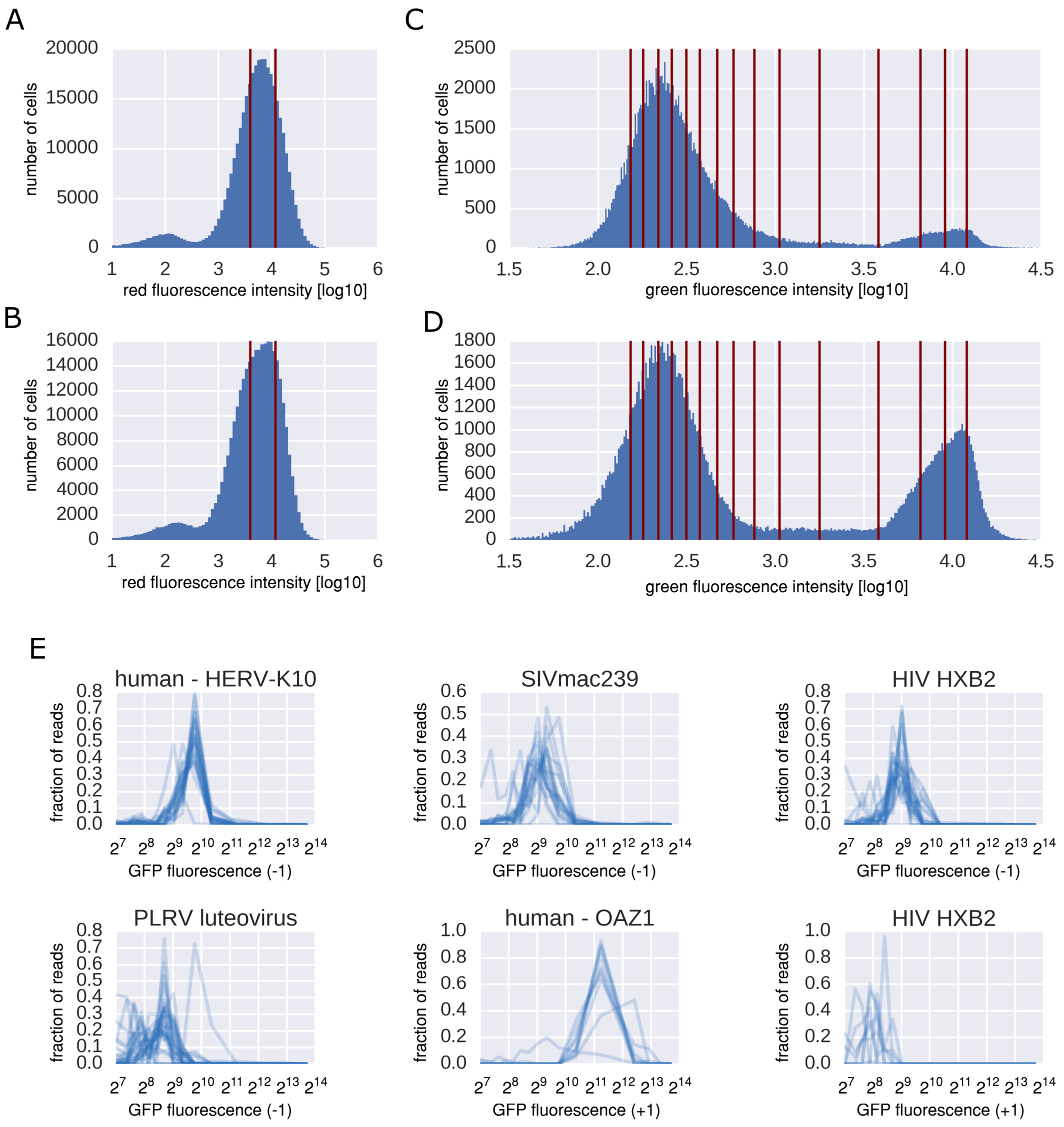


Figure S1.

AB. Histograms of red fluorescence intensity of a population of cells each carrying a -1 PRF (A) or +1 PRF (B) reporter construct from the library; red vertical lines mark the narrow range of mCherry expression used for subsequent sorting into bins of GFP expression. CD. Histograms of green fluorescence intensity of a population of cells with a narrow range of mCherry expression levels (panels AB); red lines denote the borders between the 16 bins, into which the population is subsequently sorted. E. Bin profiles (after sorting by GFP fluorescence of the -1 PRF or +1 PRF reporter construct) of barcode control variants for the indicated PRF events.

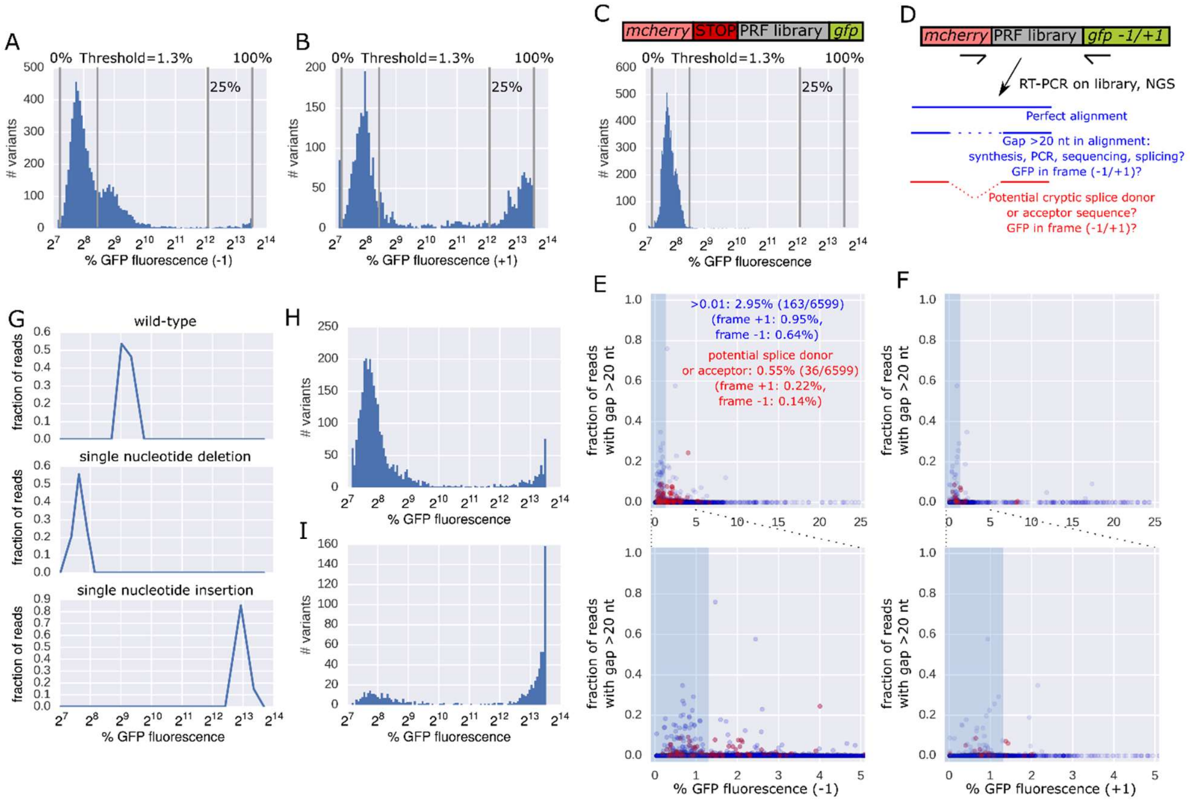


Figure S2.

AB. Distribution of mean GFP fluorescence for all variants in the -1 (A) or +1 (B) PRF reporter library. C. Distribution of percent GFP fluorescence of all measured library sequences in a reporter construct containing a stop cassette before the potential frameshift inducing sequence (the outline of the construct is shown on top). D. Outline of the experimental setup for quantifying the incidence of cryptic splicing across the entire library. EF. The fraction of sequencing reads with a gap of >20 nt (possibly resulting from an event of cryptic splice site usage) is plotted against percent GFP fluorescence (-1 frame (E) or +1 frame (F)) for all tested library variants; the shaded area denotes the range of background fluorescence. G. Bin profiles for variants of a library sequence randomly generated during synthesis and cloning; top: wild-type sequence; middle: single nucleotide deletion leading to GFP being in frame +1 and therefore no longer made into protein when -1 frameshifting occurs; bottom: single nucleotide insertion that leads to GFP being in frame with mCherry (GFP expression is not conditional on a frameshift anymore). HI. Distribution of percent GFP fluorescence of all measured variants of library sequences containing a single nucleotide deletion (H) or a single nucleotide insertion (I).

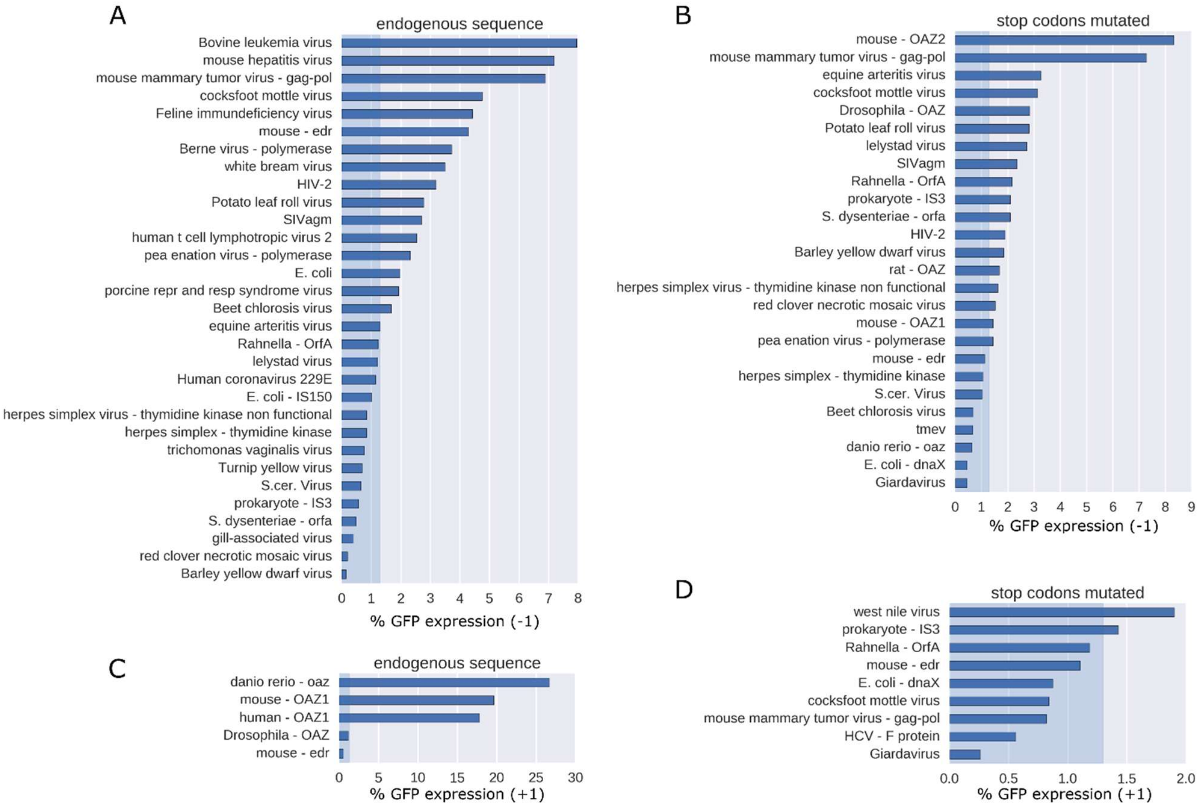


Figure S3.

A-D. Median GFP fluorescence (-1 frame, AB, or +1 frame, CD) of reported or predicted PRF variants, with (BD) or without (AC) mutating downstream stop codons, n>2 for each; the shaded area denotes the range of background fluorescence.

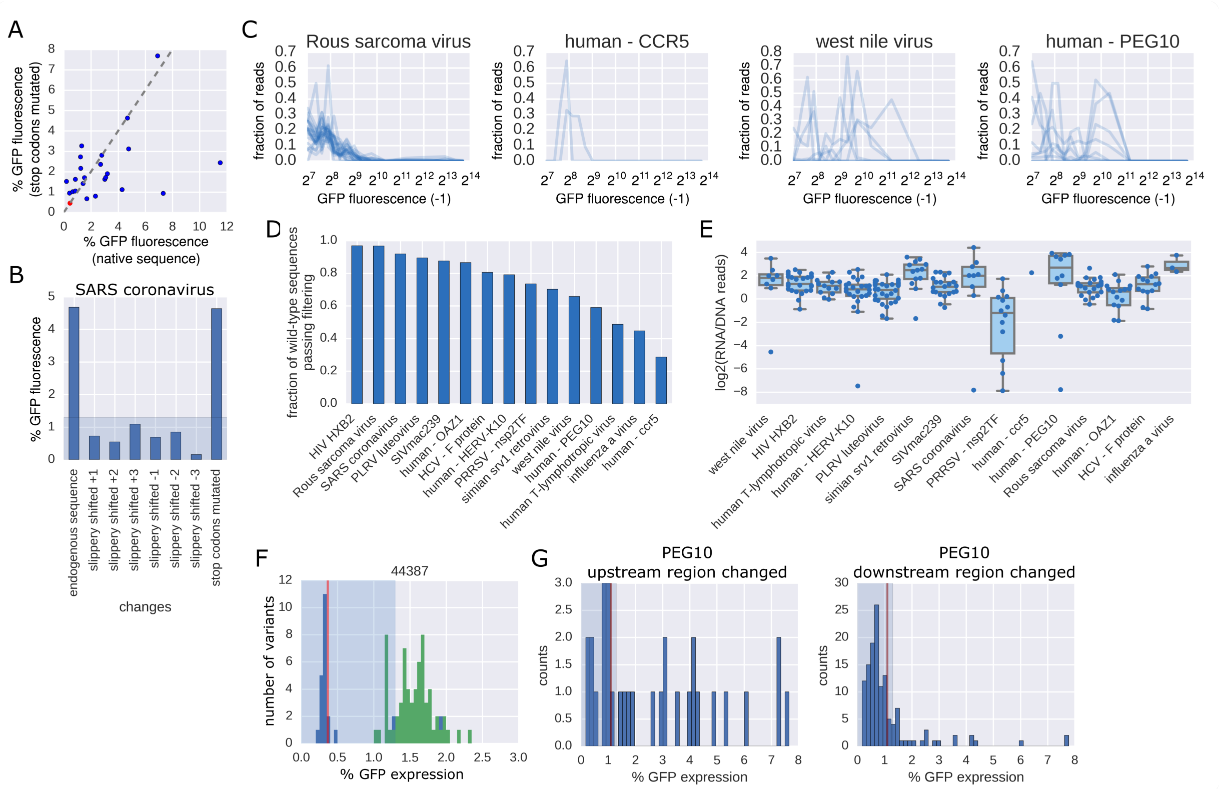


Figure S4.

A. % GFP fluorescence of library variants with all stop codons downstream of the slippery site mutated (to TGG) are plotted against % GFP fluorescence of library variants carrying the corresponding native sequence. B. % GFP fluorescence of SARS coronavirus library variants with the slippery site shifted by 1-3 nt in the 5' or 3' direction or all stop codons downstream of the slippery site mutated (to TGG). C. Bin profiles (after sorting by GFP fluorescence of the -1 PRF or +1 PRF reporter construct) of barcode control variants for the indicated PRF events. D. Fraction of library variants of wild-type sequences of the indicated PRF event passing filtering. E. RNA levels (log2(RNA/DNA reads)) for PRF reporters with the native frameshifting site (multiple barcodes, same variable region). F. Histogram of percent GFP fluorescence of all variants of a library sequence randomly generated during synthesis and cloning; the red vertical line denotes the value for the wild-type sequence, blue denotes values coming from all variants, green denotes values coming from variants containing a point mutation at position 7 after the slippery site; the shaded area denotes the range of background fluorescence. G. Distribution of all library variants based on the PEG10 PRF site with mutations exclusively in the upstream region (left), slippery site (middle) or downstream region (right); the shaded area denotes the range of background fluorescence.

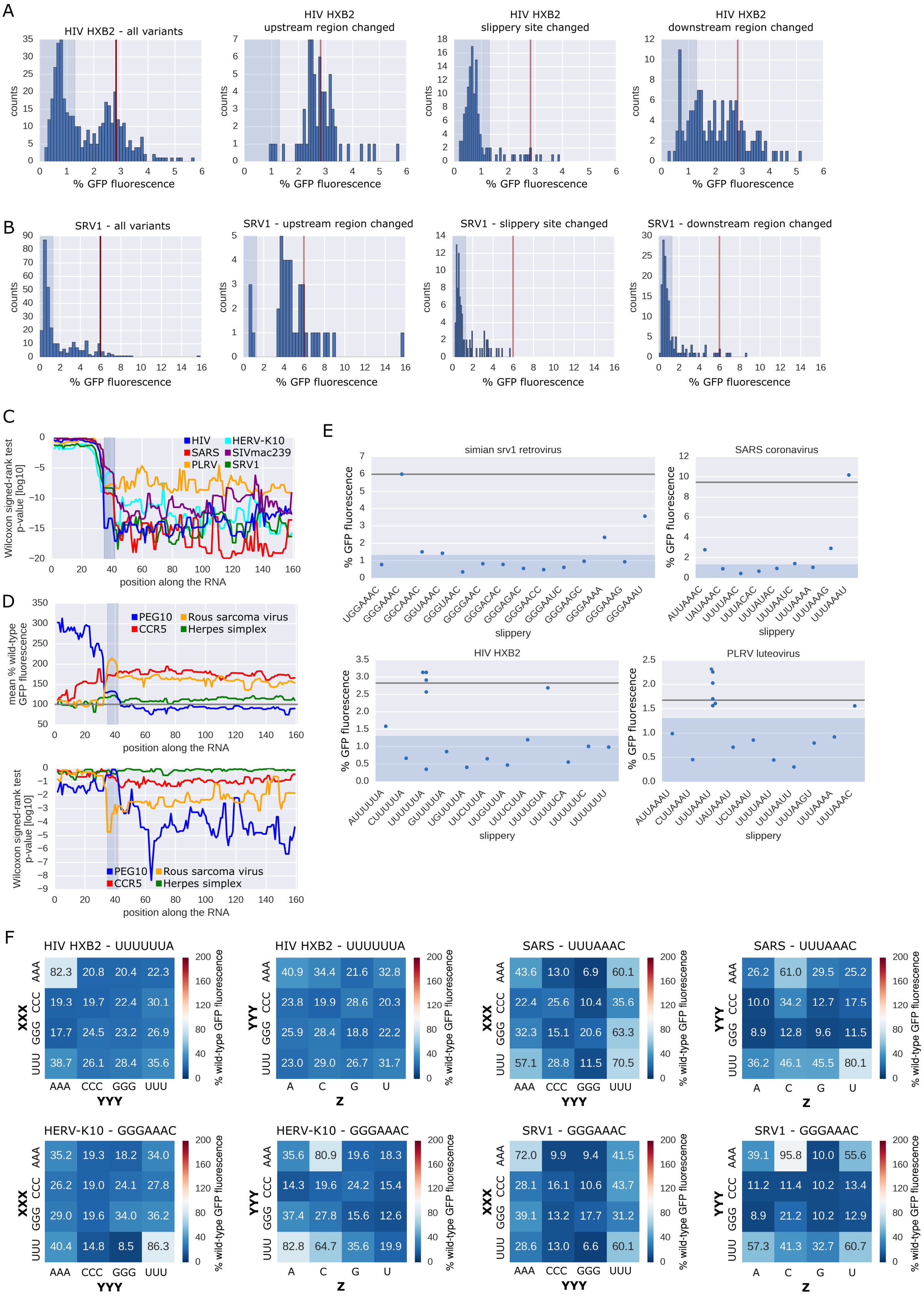

A
HIV HXB2 - all variants
HIV HXB2 upstream region changed
HIV HXB2 slippery site changed
HIV HXB2 downstream region changed
counts
% GFP fluorescence
B
SRV1 - all variants
SRV1 - upstream region changed
SRV1 - slippery site changed
SRV1 - downstream region changed
counts
% GFP fluorescence
C
Wilcoxon signed-rank test p-value [log10]
HIV
HERV-K10
SARS
SIVmac239
PLRV
SRV1
position along the RNA
D
mean % wild-type GFP fluorescence
PEG10
Rous sarcoma virus
CCR5
Herpes simplex
position along the RNA
Wilcoxon signed-rank test p-value [log10]
PEG10
Rous sarcoma virus
CCR5
Herpes simplex
position along the RNA
E
simian srv1 retrovirus
% GFP fluorescence
UGGAAAC
GGGAAAC
GGCAAAC
GGUAAAC
GGGUAAC
GGGGAAC
GGGACAC
GGGAGAC
GGGAACC
GGGAAUC
GGGAAGC
GGGAAAA
GGGAAAG
GGGAAAU
slippery
SARS coronavirus
% GFP fluorescence
AUUAAAC
UAUAAAC
UUUUAAC
UUUACAC
UUUAUAC
UUUAAUC
UUUAAAA
UUUAAAG
UUUAAAU
slippery
HIV HXB2
% GFP fluorescence
AUUUUUA
CUUUUUA
UUUUUUA
GUUUUUA
UGUUUUA
UUCUUUA
UUGUUUA
UUUCUUA
UUUUGUA
UUUUUCA
UUUUUUC
UUUUUUU
slippery
PLRV luteovirus
% GFP fluorescence
AUUAAAU
CUUAAAU
UUUAAAU
UAUAAAU
UCUAAAU
UUUUAAU
UUUAAUU
UUUAAGU
UUUAAAA
UUUAAAC
slippery
F
HIV HXB2 - UUUUUUA
XXX / YYY
AAA: 82.3 20.8 20.4 22.3
CCC: 19.3 19.7 22.4 30.1
GGG: 17.7 24.5 23.2 26.9
UUU: 38.7 26.1 28.4 35.6
AAA CCC GGG UUU
% wild-type GFP fluorescence
HIV HXB2 - UUUUUUA
YYY / Z
AAA: 40.9 34.4 21.6 32.8
CCC: 23.8 19.9 28.6 20.3
GGG: 25.9 28.4 18.8 22.2
UUU: 23.0 29.0 26.7 31.7
A C G U
% wild-type GFP fluorescence
SARS - UUUAAAC
XXX / YYY
AAA: 43.6 13.0 6.9 60.1
CCC: 22.4 25.6 10.4 35.6
GGG: 32.3 15.1 20.6 63.3
UUU: 57.1 28.8 11.5 70.5
AAA CCC GGG UUU
% wild-type GFP fluorescence
SARS - UUUAAAC
YYY / Z
AAA: 26.2 61.0 29.5 25.2
CCC: 10.0 34.2 12.7 17.5
GGG: 8.9 12.8 9.6 11.5
UUU: 36.2 46.1 45.5 80.1
A C G U
% wild-type GFP fluorescence
HERV-K10 - GGGAAAC
XXX / YYY
AAA: 35.2 19.3 18.2 34.0
CCC: 26.2 19.0 24.1 27.8
GGG: 29.0 19.6 34.0 36.2
UUU: 40.4 14.8 8.5 86.3
AAA CCC GGG UUU
% wild-type GFP fluorescence
HERV-K10 - GGGAAAC
YYY / Z
AAA: 35.6 80.9 19.6 18.3
CCC: 14.3 19.6 24.2 15.4
GGG: 37.4 27.8 15.6 12.6
UUU: 82.8 64.7 35.6 19.9
A C G U
% wild-type GFP fluorescence
SRV1 - GGGAAAC
XXX / YYY
AAA: 72.0 9.9 9.4 41.5
CCC: 28.1 16.1 10.6 43.7
GGG: 39.1 13.2 17.7 31.2
UUU: 28.6 13.0 6.6 60.1
AAA CCC GGG UUU
% wild-type GFP fluorescence
SRV1 - GGGAAAC
YYY / Z
AAA: 39.1 95.8 10.0 55.6
CCC: 11.2 11.4 10.2 13.4
GGG: 8.9 21.2 10.2 12.9
UUU: 57.3 41.3 32.7 60.7
A C G U
% wild-type GFP fluorescence

Figure S5.

AB. Distribution of all library variants based on the HIV (A) or SRV1 (B) PRF site (left) or those with mutations exclusively in the upstream region (center left), slippery site (center right) or downstream region (right); the shaded area denotes the range of background fluorescence. C. p-values (Wilcoxon signed-rank test) for the effect of a position being mutated on percent GFP fluorescence (compared to the wild-type value). D. The median percent wild-type PRF signal of variants in which the indicated position is changed (top) and the associated p-value (Wilcoxon signed rank test, bottom) are plotted for the entire length of the variable region for PRF events whose wild-type sequence did not yield a signal above threshold in our assay; gray box: slippery site. E. Percent GFP fluorescence of variants with the slippery site replaced by the indicated sequence; the horizontal line denotes the wild-type value; the shaded area denotes the range of background fluorescence. F. Heatmaps showing median percent of wild-type frameshifting conferred by a combination of slippery site elements (rows and columns) in the context of the indicated PRF event (n=2-8 for each combination).

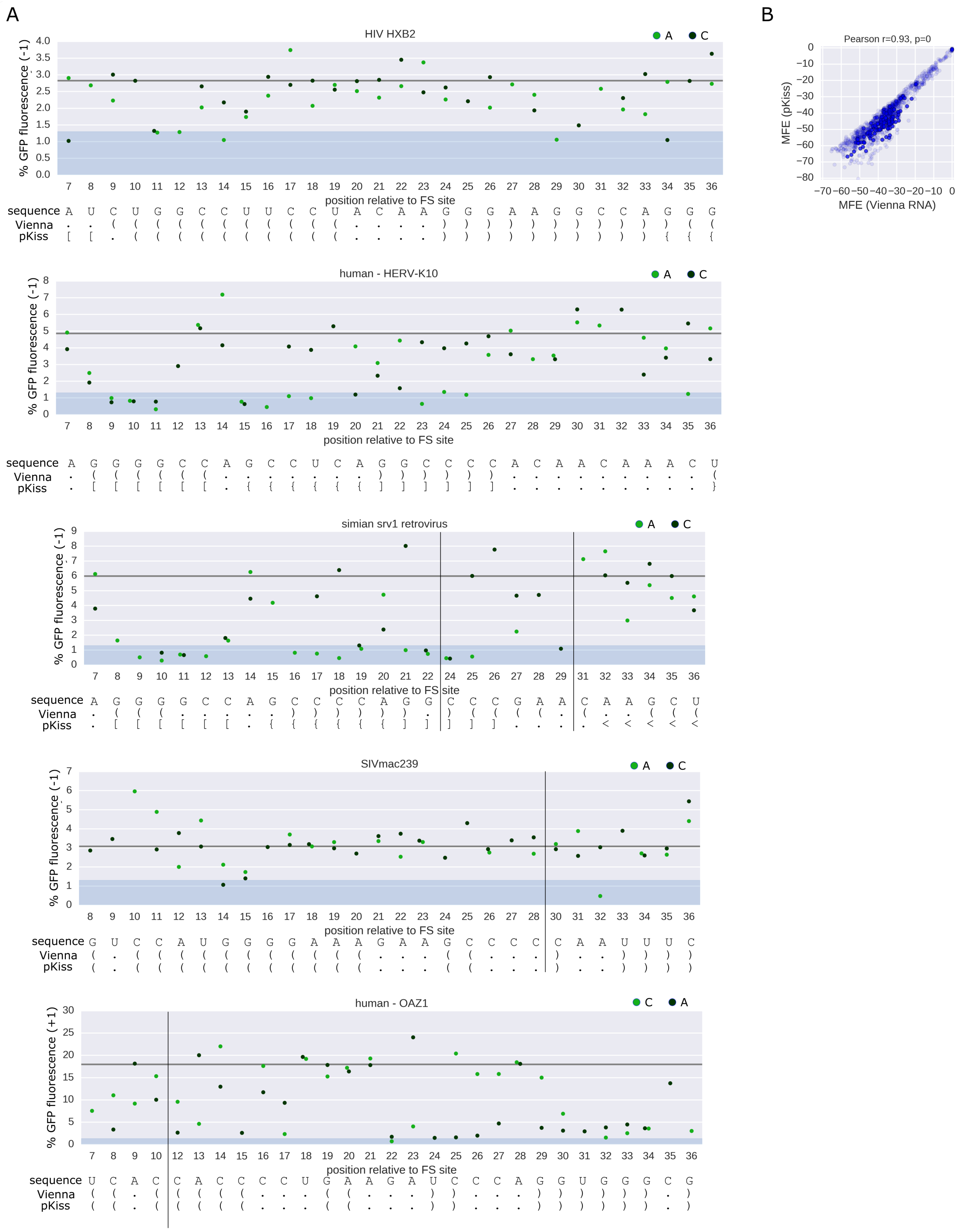


Figure S6.

A. Percent GFP fluorescence (+1 in the case of OAZ (bottom), -1 for the rest) of variants with the indicated point mutations (C or A introduced at a single position after the slippery site); the primary sequence and the predicted secondary structure (based on ViennaRNA or pKiss prediction run on the entire downstream region (120 nt)); the shaded area denotes the range of background fluorescence. B. Comparison of MFE of the downstream region (120 nt) as calculated using RNAfold (Vienna RNA package, x-axis) or pKiss (y-axis).

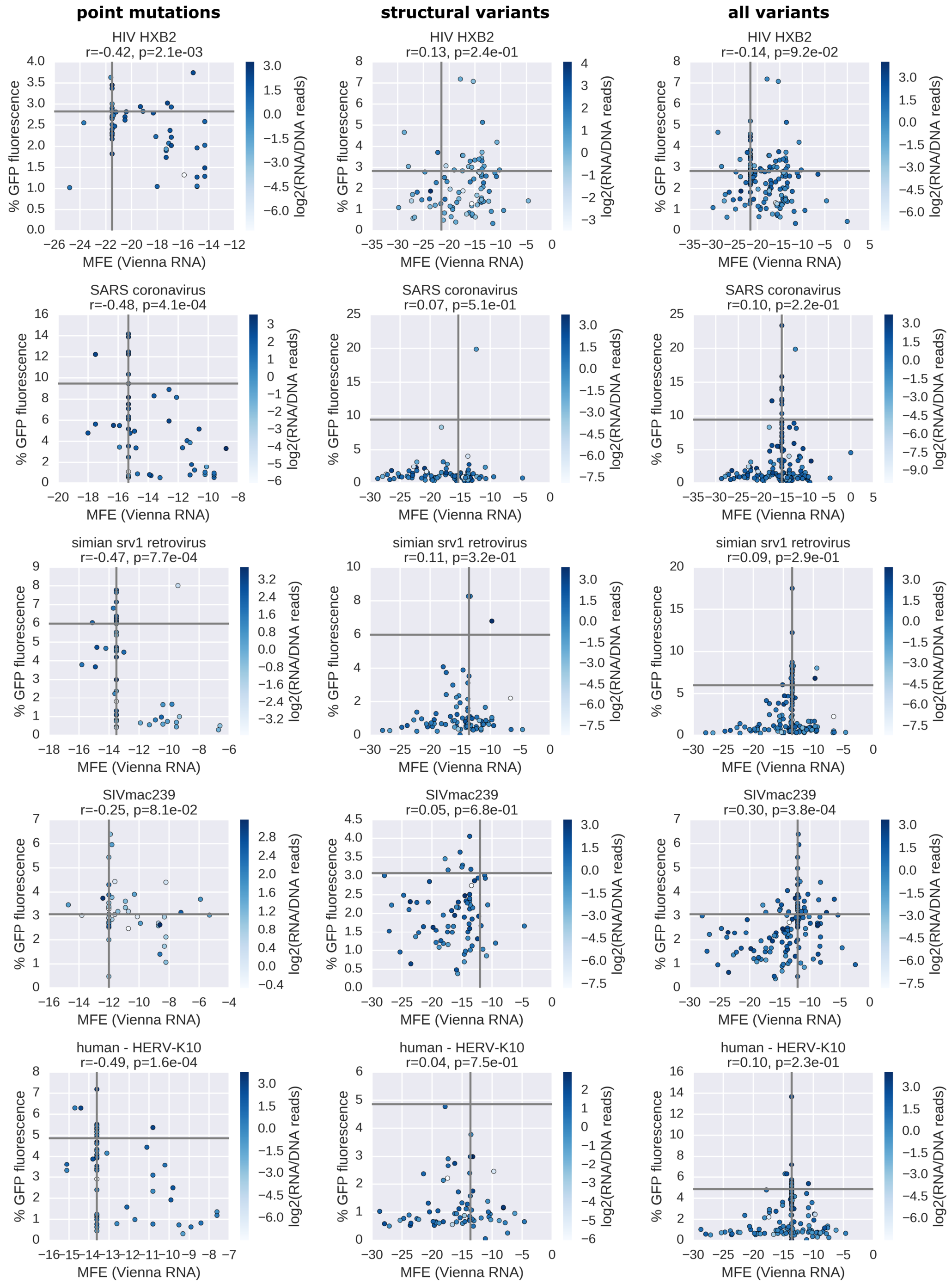


Figure S7.

Percent -1 GFP fluorescence is plotted against the MFE of the 40 nt after the slippery site (calculated using Vienna 2.0 RNAfold) for groups of variants of the indicated PRF sites; horizontal lines denote wild-type percent GFP fluorescence, vertical lines denote the MFE of the wild-type sequence, shades of blue denote RNA levels (log2(RNA/DNA reads)).

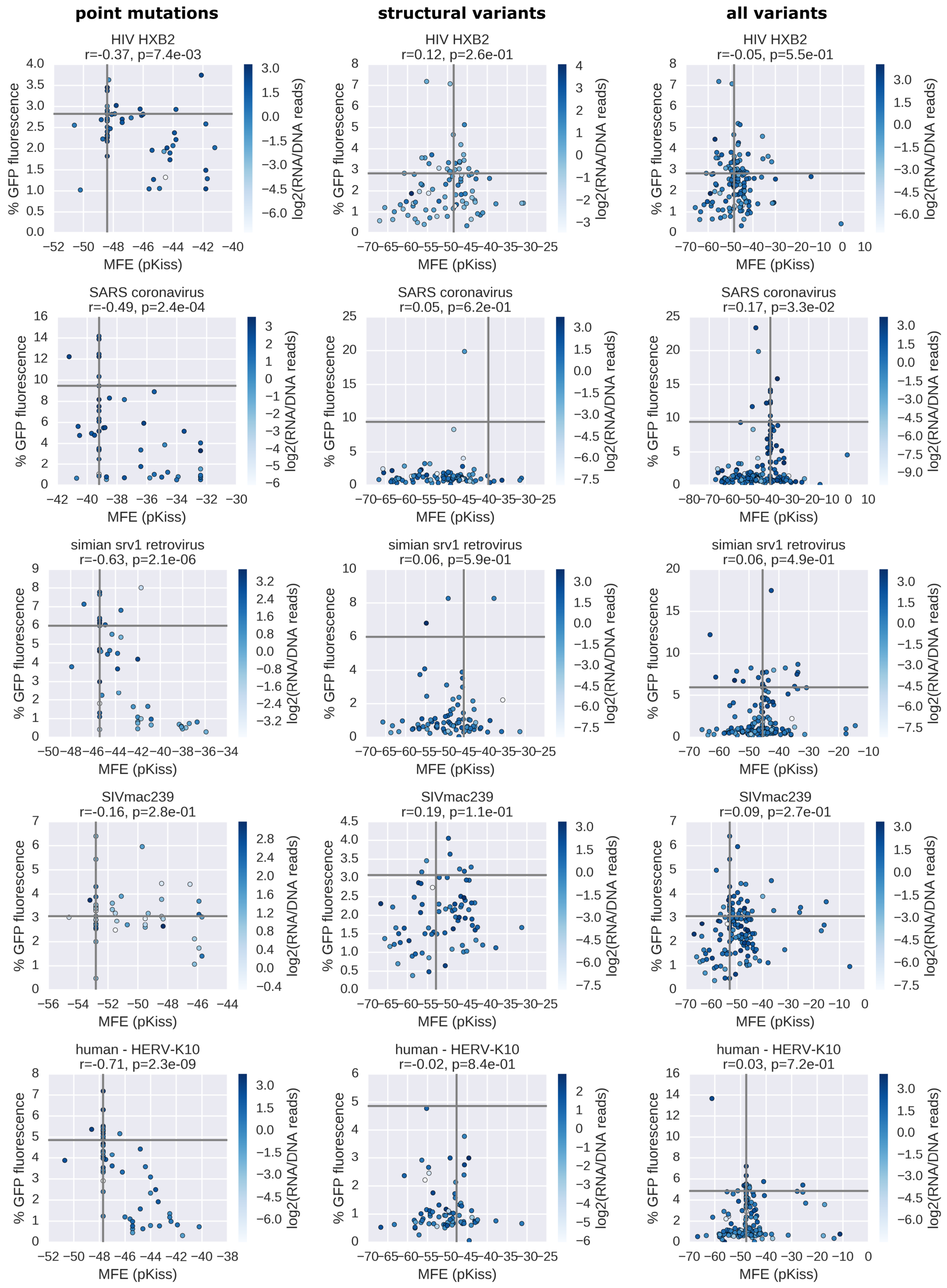


Figure S8.

Percent -1 GFP fluorescence is plotted against the MFE of the 120 nt after the slippery site (calculated using pKiss) for groups of variants of the indicated PRF sites; horizontal lines denote wild-type percent GFP fluorescence, vertical lines denote the MFE of the wild-type sequence, shades of blue denote RNA levels (log2(RNA/DNA reads)).

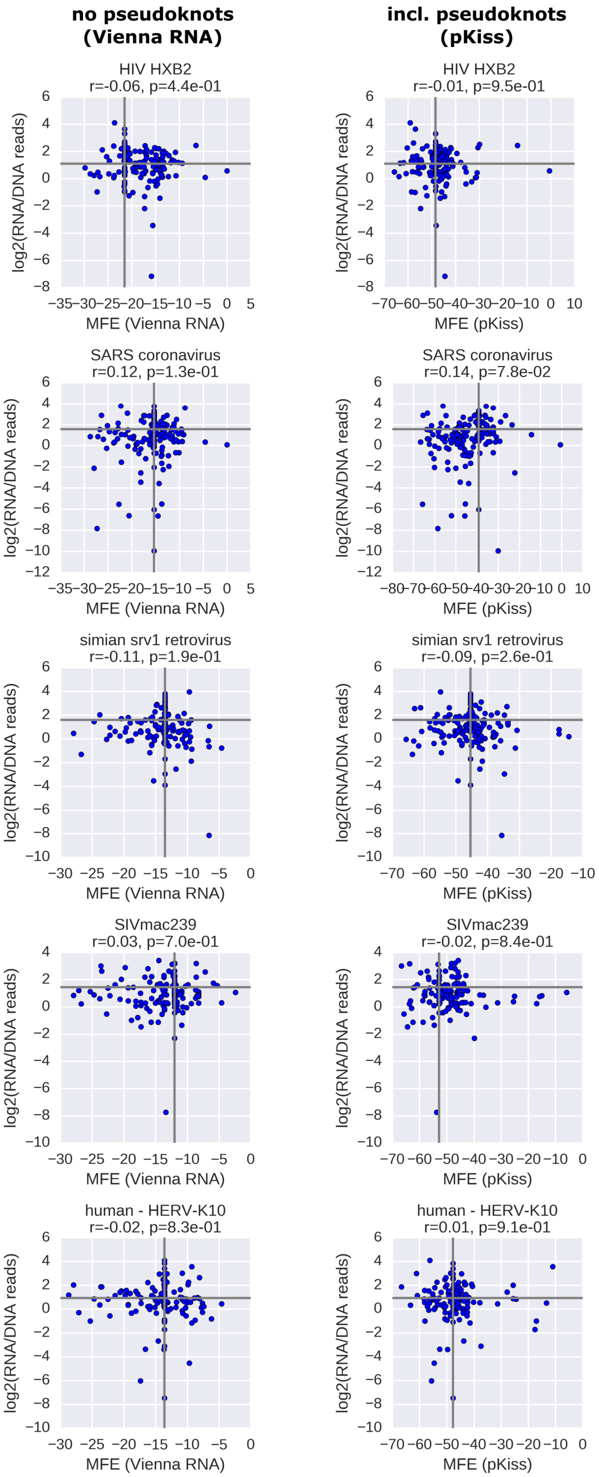


Figure S9.

RNA levels (log2(RNA/DNA reads)) are plotted either against the MFE of the 40 nt after the slippery site (calculated using Vienna 2.0 RNAfold) or against the MFE of the 120 nt after the slippery site (calculated using pKiss) for groups of variants of the indicated PRF sites; horizontal lines denote wild-type RNA levels, vertical lines denote the MFE of the wild-type sequence.

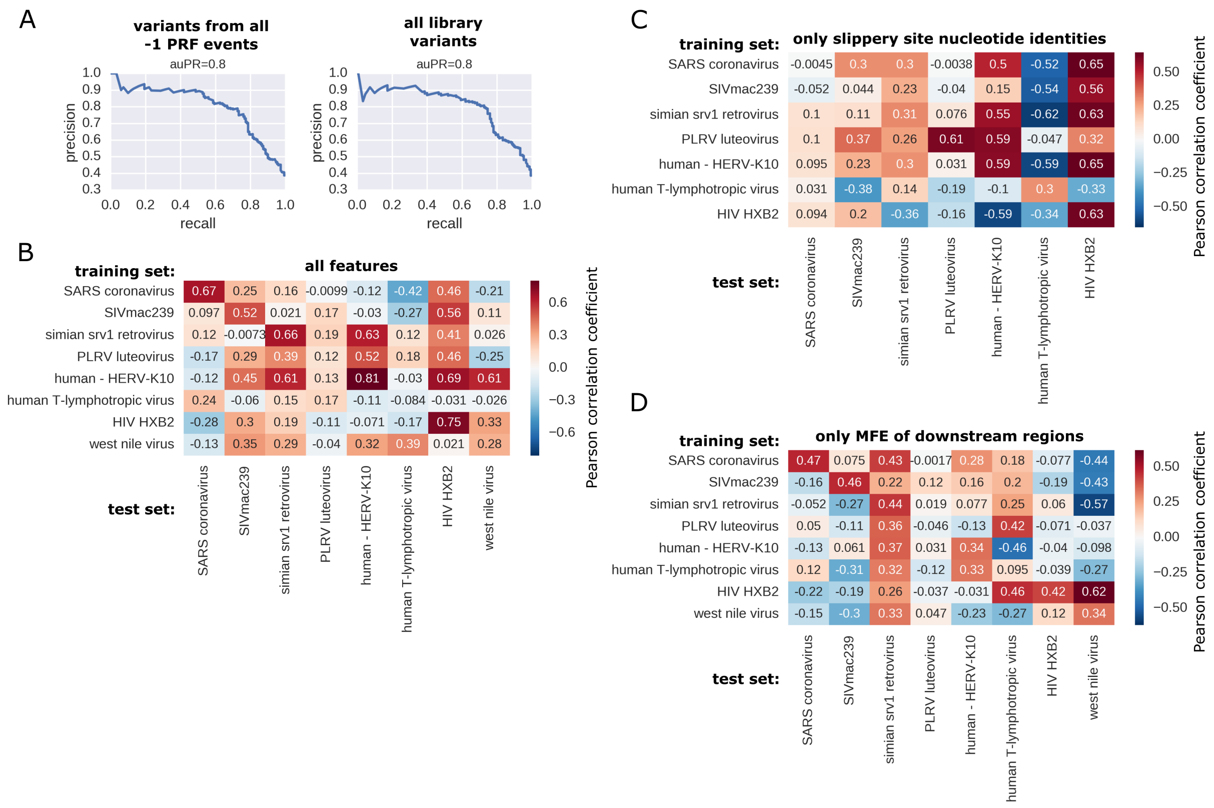


Figure S10.

A. Precision recall curves for the XGBoost classifier predicting frameshifting potential (-1 PRF signal above threshold (=1.3%)). B-D. Prediction scores (Pearson correlation coefficients) on held-out test data (20%) for the indicated PRF event (columns) using all feature sets combined (B) the identity of slippery site positions (C) or MFE of downstream regions (D) as features and training the model on the training data (80%)

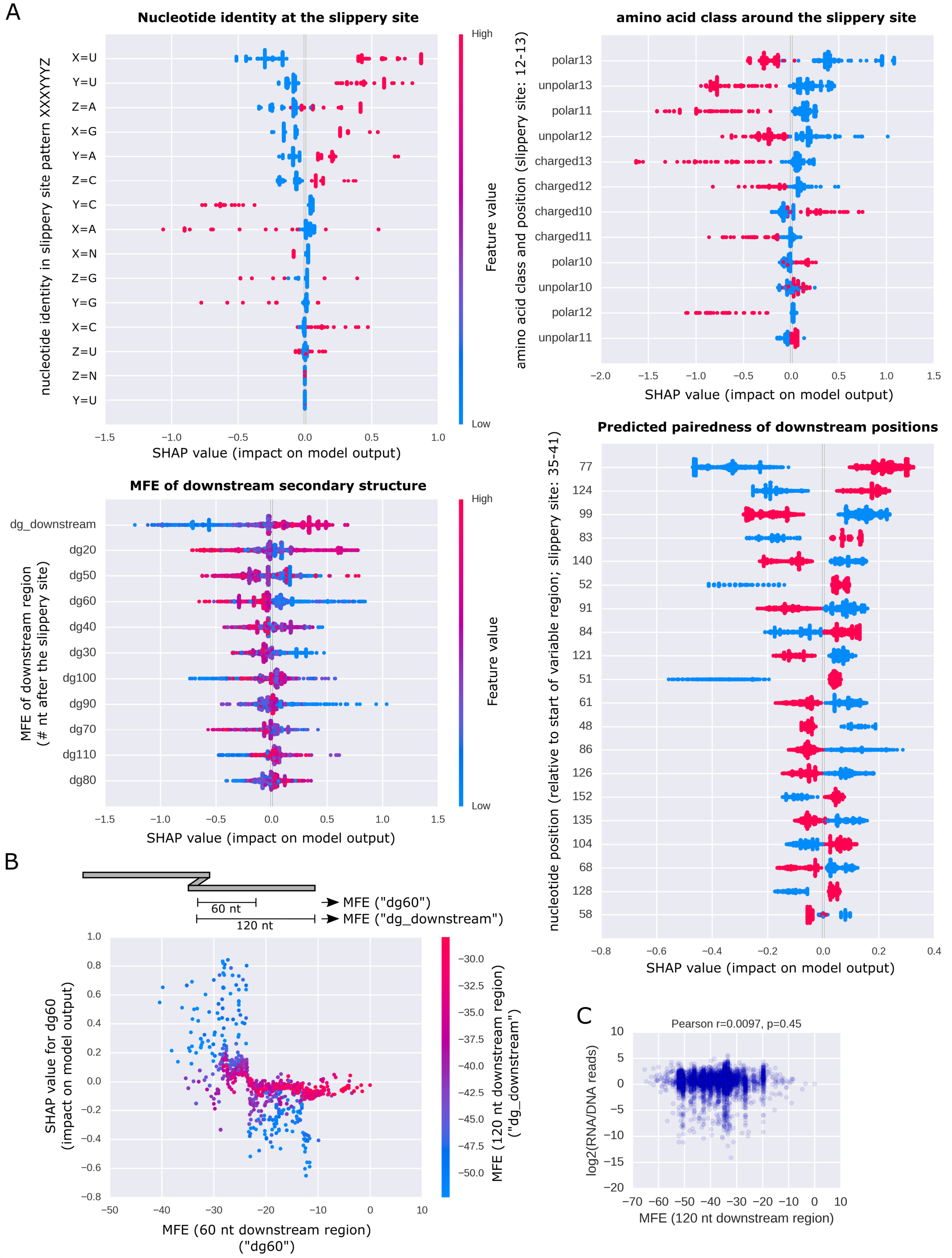

A
Nucleotide identity at the slippery site
nucleotide identity in slippery site pattern XXXYYYZ
X=U
Y=U
Z=A
X=G
Y=A
Z=C
Y=C
X=A
X=N
Z=G
Y=G
X=C
Z=U
Z=N
Y=U
SHAP value (impact on model output)
Feature value
High
Low
amino acid class around the slippery site
amino acid class and position (slippery site: 12-13)
polar13
unpolar13
polar11
unpolar12
charged13
charged12
charged10
charged11
polar10
unpolar10
polar12
unpolar11
SHAP value (impact on model output)
MFE of downstream secondary structure
MFE of downstream region (# nt after the slippery site)
dg_downstream
dg20
dg50
dg60
dg40
dg30
dg100
dg90
dg70
dg110
dg80
SHAP value (impact on model output)
Predicted pairedness of downstream positions
nucleotide position (relative to start of variable region; slippery site: 35-41)
77
124
99
83
140
52
91
84
121
51
61
48
86
126
152
135
104
68
128
58
SHAP value (impact on model output)
B
60 nt
120 nt
MFE ("dg60")
MFE ("dg_downstream")
SHAP value for dg60 (impact on model output)
MFE (60 nt downstream region) ("dg60")
MFE (120 nt downstream region) ("dg_downstream")
C
Pearson r=0.0097, p=0.45
log2(RNA/DNA reads)
MFE (120 nt downstream region)

Figure S11.

A. Effects (as determined using SHAP) of slippery site identity (in the canonical pattern XXXYYYZ), amino acid properties (numbers indicated position relative to the start of the variable region (not including the barcode); slippery site=12-13), pairedness (numbers indicated position relative to the start of the variable region (not including the barcode); slippery site=35-41) and MFE (numbers indicate the size of the downstream region for which the MFE was calculated using RNAfold (Vienna 2.0); “downstream”: 120 nt) features on the model prediction (ranked by their importance for the prediction) for a classifier built on the indicated feature sets. The color denotes the feature value and the position along the x-axis denotes the impact on model output, for each item in the training set. B. The impact (y-axis) of MFE of the first 60 nt after the slippery site (x-axis) dependent on the MFE calculated on the full downstream region (120 nt after the slippery site; color scale) on the model prediction (as determined using SHAP) is plotted against the feature values for each item in the training set. C. Comparison of MFE of the downstream region (120 nt) as calculated using RNAfold (Vienna RNA package, x-axis) and RNA levels (log2(RNA/DNA reads)) for all library variants used for training and testing the models.

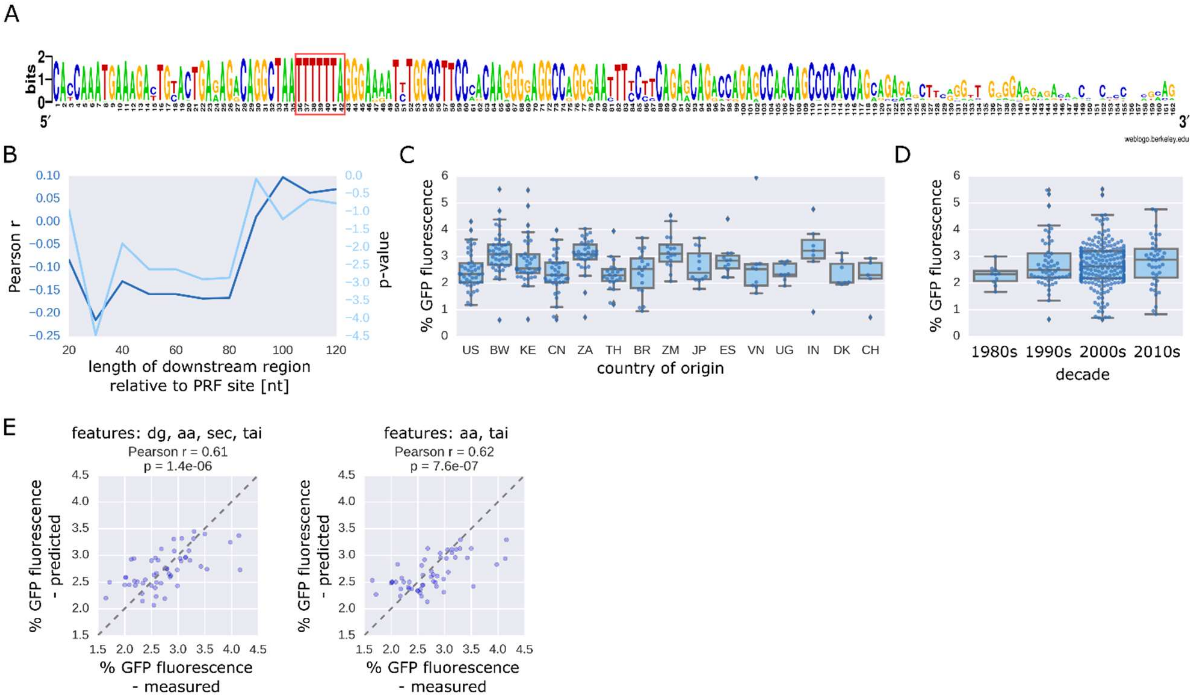


Figure S12.

A. Sequence logo created from multiple alignment of all tested HIV variants. B. Pearson correlation coefficient (dark blue) and the associated p-value (light blue) between minimum free energy and percent GFP fluorescence of HIV variants are shown for secondary structure predictions based on the indicated length of the downstream region. CD. Boxplot showing percent GFP fluorescence of HIV gag-pol PRF variants coming from the indicated country of origin (C) or decade (D). E. Predicted vs measured percent GFP fluorescence for 20% of the HIV variants (held-out test set) with the full feature set except for the identity of slippery site positions (left) or after removing MFE features (middle) or all secondary structure derived features (right) features and trained on both the other 80% of the HIV variants and the designed HIV variants combined.
